# Bacterial hopanoids: a newly identified potent inducer of monocyte to macrophage differentiation

**DOI:** 10.1101/2021.03.20.436240

**Authors:** Anindita Bhattacharya, Arnab Ghosh, Suman Mallik, Subhasis Mandal, Prosenjit Sen

## Abstract

Monocyte to macrophage differentiation is an extremely essential cellular and immunological process aimed to combat the assault of foreign pathogens as well as this process has immense importance in the maintenance of physiological homeostasis. Differentiated macrophage, serving as specialized phagocytes is an indispensable component of innate immunity. Besides this, being a well-documented antigen-presenting cell (APC) macrophages also function as a key regulator of both humoral and cell-mediated immunity: the two integral components of adaptive immunity. Bacterial hopanoids are a primitive analogous of eukaryotic sterols that are established as a prominent immunomodulator. In this study, we have demonstrated the isolation protocol of neutral lipid fractions from different Gram-negative bacteria, and adopting several analytical approaches we proposed that the extracted lipid fractions may contain hopanoid as an active component. We have delved deeper into the biological effect of these probable hopanoids like compounds. Here, by considering the several structural and functional attributes like altered phenotype, expression of macrophage-specific markers, increased intracellular organelles, acquirement of enhanced phagocytic and inflammatory property, induction of autophagy, etc we have established hopanoids as a potential inducer of monocyte to macrophage differentiation. Thus, our study has unraveled a new immunomodulator present in Gram-negative bacteria and would undoubtedly help to understand the intricacies of host-pathogen interaction in a better and conclusive manner.

## Introduction

Blood circulating monocytes and tissue-resident macrophages comprise the principal components of the mononuclear phagocytic system and are therefore considered as the chief constituents of the innate immune response[1]. Following their origin in the bone marrow, the promonocytes enter the blood circulation and further differentiate into mature monocytes[2][3]. In response to certain cellular triggers, namely pro-inflammatory signals, metabolic or immune stimuli, etc. circulating monocytes infiltrate peripheral extravascular tissues and undergo concomitant differentiation into either macrophages or dendritic cells[4]. As vigilant sentinels, macrophages are entrusted with the vital responsibility of protecting the system against the onslaught of invading foreign pathogens[5]. Macrophages occupy specific territories in organs where they contribute to host defense, development, tissue-remodeling and repair[6]. The phenomenon of monocyte to macrophage differentiation is firmly regulated and a slight deviation from normalcy can result in severe life-threatening conditions.

For a long time different research groups, working with monocytes have chosen the microscope-based study as a very authentic mode for analyzing monocyte morphology. Circulating Monocytes havinga diameter of 10-15μm acquires an amoeboid appearance[7]. They have a granulated cytoplasm[8] and almost half of the cellular volume is occupied by the eccentrically placed reniformnucleus [9]. They also have a lesser number of filopodia as well as membrane-bound cytosolic organelles. On the other hand, differentiating cells exhibit an enlargedsize, numerouscytoplasmic projections, enhanced spreading ability, autophagy induction as well as altered surface protein expression profile[10][11][12,13]. As a crucial regulator of the innate immune system differentiated macrophages engulf foreign pathogens like fungi, bacteria, and protozoans[1]. Through the production of an array of inflammatory mediators like cytokines, interleukins, ROS, NO the differentiated macrophages also destroy the dead and damaged cells and aid in the process of tissue repair[14][15][16][17][17]. Thus monocyte/macrophage elicits an explicit immune response against both intra and extracellular pathogens.Apart from a multitude of well-established chemical stimuli like GM CSF, MCSF, PMA, LPS, etc many biologically active compounds includingnatural products are known to induce monocyte to macrophage differentiation. Different hormones are also reported to induce differentiation. Although monocyte to macrophage differentiation is indispensable for our immune system, recent scientific developments strongly advocate the involvement of monocytes and monocyte-derived macrophages in several diseases like systemic sclerosis, Adult-Onset Still’s Disease (AOSD), primary biliary cholangitis, rheumatoid arthritis, etc. Therefore the process of monocyte to macrophage differentiation has profound significance on cellular physiology and mandates an in-depth study[18][19][20][21].

Hopanoids are the primitive analogue of sterol present predominantly in bacteria. They are steroid-like compoundsrepresentative of a populous group of pentacyclic triterpenoids:a class of chemical compounds composed of three terpene units with the molecular formula C_30_H_48_. They may also be classified as a compound consisting of six isoprene units. Although triterpenoids are almost ubiquitously present in a diverse group of organisms; hopanoids are found in bacteria, lichens, bryophytes, ferns, and fungi[22].Hopanoidsare hypothesized to be surrogates of eukaryotic sterols primarily because of similarities in their biosynthesis schemes and structure[23][24][25][26][27]. Hopanoidsare found to localize in the cytoplasmand outer membranes of certain bacteria[28,29][30]. Results from *in-vitro* experiments have shown that they can condense artificial membranes and enhance the viscosity of liposomes[31][32]. Through a genetic approach, Welander et al have shown that loss of hopanoid production in *Rhodopseudomonaspalustris*leads to enhanced sensitivity to acidic and alkaline conditions, bilesalts, and certain antibiotics[33]. Although the deletion of genes involved in hopanoid synthesis results in nonlethal phenotype in bacteria, hopanoid-deficientmutants have been shown to exhibit increased sensitivity to antibiotics and detergents as well as they are more susceptible to stresses, including variation in pH, temperature, and osmotic pressure[33][34][35][36][37]. Peter et al have reported that hopanoids interact with lipid A like cholesterol and sphingolipids to order membranes[38].

Although the effect of bacterial hopanoids on host physiology and the immune system is not characterized yet, few reports are suggesting the immunomodulatory activity of other triterpenoids from various sources. Eight cycloartanes isolated from *Astragalusmelanophrurius* (Fabaceae) are reported to stimulate humanlymphocyte proliferation[39]. The mixtureof triterpenes from *Quillajasaponaria* (Rosaceae) stimulates the production of inflammatory cytokines like IL-1, IL-6, etc along with activation of antigen-presenting cells (APCs) for the elicitation of the immune response[40]. Khajuria et al isolated oleanolic acidand echinocystic acid from *Luffacylindrica* (Cucurbitaceae) and reported their immune-modulatoryactivity.Both compounds enhanced the phagocytic index and imparted stimulatory effects on macrophages, and thereby triggers both humoral and cell-mediated immune responses[41]. On the other hand, in the recent past, Qu et al have shown anticancer, antioxidant, and liver-protective effects of triterpenes from *Ganodermalucidum*[42].

Through this study,we investigated the novel effect of bacterial hopanoids on the host immune system.Although the immunomodulatory activity of LPS present in Gram-negative bacteria is known for years, the contribution of other bacterial components is not delved in detail yet.The structural similarities between bacterial hopanoids with those of sterol based immunomodulator led us to hypothesize that hopanoids may be a potent inducer of monocyte to macrophage differentiation. Like LPS, hopanoids can come to the immediate proximity of the circulating monocytes as it is an integrated component of the bacterial outer cell membrane, otherwise, there lies some possibility of the bacterial hopanoids in the circulation originated from the lysed bacteria. In this manuscript, we have established the immunomodulatory potential of bacterial hopanoids by demonstrating the morphological, physiological as well as functional attributes of differentiating monocytes in response to bacterial hopanoids treatment. Identification of this new immunomodulator will undoubtedly aid in the study of host-pathogen interaction as well as it will be useful for the design of new antibacterial therapeutics.

## Material and Methods

### Cell culture

Monocytic leukemia cell line (THP-1) of ATCC origin was obtained from Dr. Amitabho Sengupta, IICB. THP-1 was maintained and stimulated in RPMI-1640 medium supplemented with 10% heat-inactivated FBS and 5% Penstrep (Invitrogen and Himedia) by keeping in an incubator fixed at 37°C and equipped with 5% CO_2_ supply. For inducing monocyte to macrophage differentiation THP-1 cells were treated with PMA (50ng/ml) or LPS (100μg/ml) and seeded either on a confocal dish (for microscopy) or on a commercially available adhesion compatible dish and kept in the incubator for requisite time point.

### Western blot

According to the standard protocol, Western blot was performed. Cells were lysed with Laemmeli buffer. Lysates were loaded and separated in SDS PAGE. Then proteins were transferred to PVDF membranes (Millipore), incubated with primary and secondary antibodies and bands were detected. Pre-stained molecular markers (Biorad) were used to estimate the molecular weight of samples. The protein expression level was quantified with Image J software (NIH).

### RT-PCR

Total RNA isolation was done using Trizol (Life-technologies) according to the manufacturer’s instruction. Reverse transcription was done with life technologies kit. β-actin was used as an internal control. Subsequently desired gene was PCR amplified.

### Phagocytosis assay

Carboxylated latex fluorescent beads of 200nm (Invitrogen) diameter were addedto the cells. Two hours later the cells were washed with PBS to remove the floating beads. RFP tagged *E*.*coli* culture was added to the cells. 30 minutes later the cells were washed with PBS to remove the floating bacteria. Fluorescence Z-stack images of the cells were acquired. Maximum intensity projection image from the previously acquired Z-stacks was obtained to count the phagocytic cell percentage. The fluorescence intensity of z-projected images was scaled such that the background intensity becomes 1. The cells with average fluorescence intensity more than 10% of the background were considered as phagocytic.

### NO measurement

Griess reagent was incubated with spent medium and absorbance was measured at 540nm.

### ROS measurement

25µM DCFDA(Sigma) was incubated with cells for 30 minutes. Next, the cells were lysed with DMSO and the fluorescence was measured at Ex485 nm/Em535 nm.

### Immunostaining

Cells were fixed with 4% paraformaldehyde in PBS for 10 min, permeabilized with 0.1% Triton x 100 for 2 mins (in case of LC3 staining), washed with PBS, blocked with 5% BSA for an hour. Primary antibody (Abcam/CST) incubation was done overnight, washed with PBS and then incubated with Alexa tagged secondary antibody (CST) for 45 minutes, finally washed with PBS, counterstained with Hoechst, and imaged. Cells were imaged with the sCMOS camera (Orca Flash 4.0, Hamamatsu) on an inverted fluorescence microscope from Carl Zeiss (Axio-observer Z1) and a confocal microscope (Carl Zeiss LSM880).Image processing was done with Image J.

### CYTO-ID staining

1μl of Cyto-ID Green Detection Reagent (Enzo-lifesciences kit) was incubated with 10^5^ to 10^6^ cells/ml for 30 min under standard tissue culture conditions at 37 °C, 5% CO_2_ in the dark.At the end of the staining procedure, the excess dye was washed with PBS and imaged at 488nm excitation filter.

### DQ™-Red BSA staining

DQ™-Red BSA (10 μg/ml) was added to cells and incubated for 15 minutes at 37 °C. Then the cells were washed with PBS and imaged at 590 nm excitation filter and 620 nm emission filter.

### Bodipy Staining

Bodipy (1 mg/ml stock) was added to cells at a final concentration of 1μg/ml and incubated for 15 minutes at 37 °C. Then the cells were washed with PBS and imaged at 480 nmexcitation filter

### Lysotracker staining

Lyso Tracker™ Red DND-99 (Life Technologies) was added to the cells at a final concentration of 25nM and incubated for 30 minutes at 37° c. Then the cells were washed with PBS and imaged at 577 nmexcitation filter.

### Mitotracker staining

Mito Tracker™ Dyes (Life Technologies) was added to the cells at a final concentration of 10 nM and incubated for 30 minutes at 37° c. Then the cells were washed with PBS and imaged at 579 nmexcitation filter

### Hopanoid isolation

60mg of dried biomass was dissolved in 20ml solution containing DCM, methanol and water in a ratio of 1:2:0.9 followed by vortexing for 20 minutes. Then equal volume of 1:1 DCM and water mix was added to it followed by vortexing for 20 minutes. Next, the solution was centrifuged at 1500 RPM for 30 minutes and the lower lipid layer was collected. Then, the whole solution was dried over sodium sulfate(Water was removed) and solvent was removed on a rotary evaporator and resultant compound was purified by column chromatography. Column was packed by silica gel (60–120 mesh size)and gradually eluted with hexane, 30% ethyl acetate-hexane(v/v), and ultimately 5%CHCl_3_-Methanol. Each fraction was evaporated to dryness by rotary evaporator. Individual fraction further purified by plate thin layer column chromatography. Plate TLC was made with Silica Gel GF254. Fine separated compound was qualitatively identified by MALDI-TOF-MS and TLC. 2,3-Dihydroxybenzoic acid was used as matrix in MALDI experiment. Normal TLC was done with TLC Silica gel 60 F_254_.TLC plate was charredon hot plate by Ninhydrin, Copper acetate, and ammonium molybdate to mark the compound. Unsaturation of hopanoids were detected by iodination followed by MALDI-TOF analysis.

### MTT staining

Cells were washed with 1X PBS and incubated with MTT (Sigma) solution (1mg/ml) for 4 hours at 37°C. Subsequently the cells were lysed with DMSO and absorbance at 570 nm was measured. From the absorbance value, cell viability was calculatedusing the following formula(%) cell viability = (the absorbance of treated cells/absorbance of control cells) x 100.

### Statistical analysis

All the techniques applied here are representative of at least three (n≥3) independent experiments. The data presented here are as mean ±S.E of the mean and the differences are considered to be statistically significant at p < 0.05 using the student’s t-test. Graphpad prism was used for statistical analysis.

## Results

### Chemical characterization of the probable hopanoids in the neutral lipids of Gram-negative bacteria

*Rhodopseudomonaspalustris*is considered a rich source of different hopanoids.Wu et al and Sessions et al have successfully isolated 2 Methyl-diplopterol, 2 Methyl bacteriohopanetetrol (2Me-BHT) and their unmethylated species (diplopterol and BHT) from *Rhodopseudomonaspalustris*[43,44].Following the reported isolation and purification protocol for neutral lipid, we have adopted a combined approach of column chromatography and preparatory TLC to fractionate the bacterial total neutral lipid. Finally, the purity of each fraction was verified by Plate TLC analysis(Fig. 2Ai, Bi, Ci). Eventually,by MALDI-TOF analysis, we verified the mass of isolated compounds to be similar to that of different hopanoids eluted in the respective fraction(Fig. 1Aii, Bii, Cii).Thus we speculated that compound 1 (Diploptene), compound 2 and compound 3 (2 methyl diploterol) may be present in 1^st^ fraction (eluted in hexane), 2^nd^ fraction (eluted in 30% Ethyl acetate-Hexane), and 3^rd^ fraction (eluted in 5% Chloroform-Methanol) respectively. To further establish the identity of the isolated lipids we have performed iodination followed by MALDI-TOF analysis to detect the unsaturation in the lipids. Only fraction 1 has shown a shift of peak in MALDI TOF analysis comparable with the molecular weight of one equivalent of iodine confirming the presence of only one unsaturation in the analyte present in fraction 1, further strengthening our proposition i.e. the identity of compound 1 (Diploptene) to be present in fraction 1(Fig. 2Aiii). Next, we sought to extend our study with other bacterial species like some of the well-known Gram-negative and Gram-positive bacteria like *Escherichia coli, Staphylococcus aureus* and *Bacillus subtilis*. Since there is no pre-existing literature reporting the detailed analysis of bacterial hopanoids from the above-mentioned species, in this manuscript we applied the protocol for isolation and characterization of neutral lipids from those strains similar to that of *Rhodopseudomonaspalustris*.MALDI-TOF basedmolecular weight analysis of respective neutral lipids fraction isolated from *Escherichia coli* reveals the presence of compounds having a molecular weight of 410, 428 & 442 in the 1^st^ fraction, 2^nd^ fraction and 3^rd^ fraction respectively(Fig. 2D-F).Thus we are speculating that *Escherichia coli* (common Gram-negative bacteria) may also contain hopanoids similar to those reported for *Rhodopseudomonaspalustris*.Similar analysis with the neutral lipid isolated from Gram-positive bacteria like *Staphylococcusaureus* and *Bacillus subtilis* discloses the absence of a hopanoid type compound within these strains(Fig. S2).LPS is a designated immunomodulator present in Gram-negative bacteria known to induce monocyte to macrophage differentiation. Thus the presence of trace quantity of LPS may lead to erroneous conclusions. Carbohydrate and phosphate moiety are the two integral components of LPS. So to check whether our lipid fractions are free of LPS or not, we have performed different biochemical assays likeFehling’s test, Benedict’s test and Seliwanoff’stestto detect the presence of carbohydrate and Phosphate NMR for the identification of phosphate in the lipid preparation. The negative result in all the assays has led us to conclude that isolated lipid fractions exclude the possibility of having any LPS contamination.

**Figure 1.**
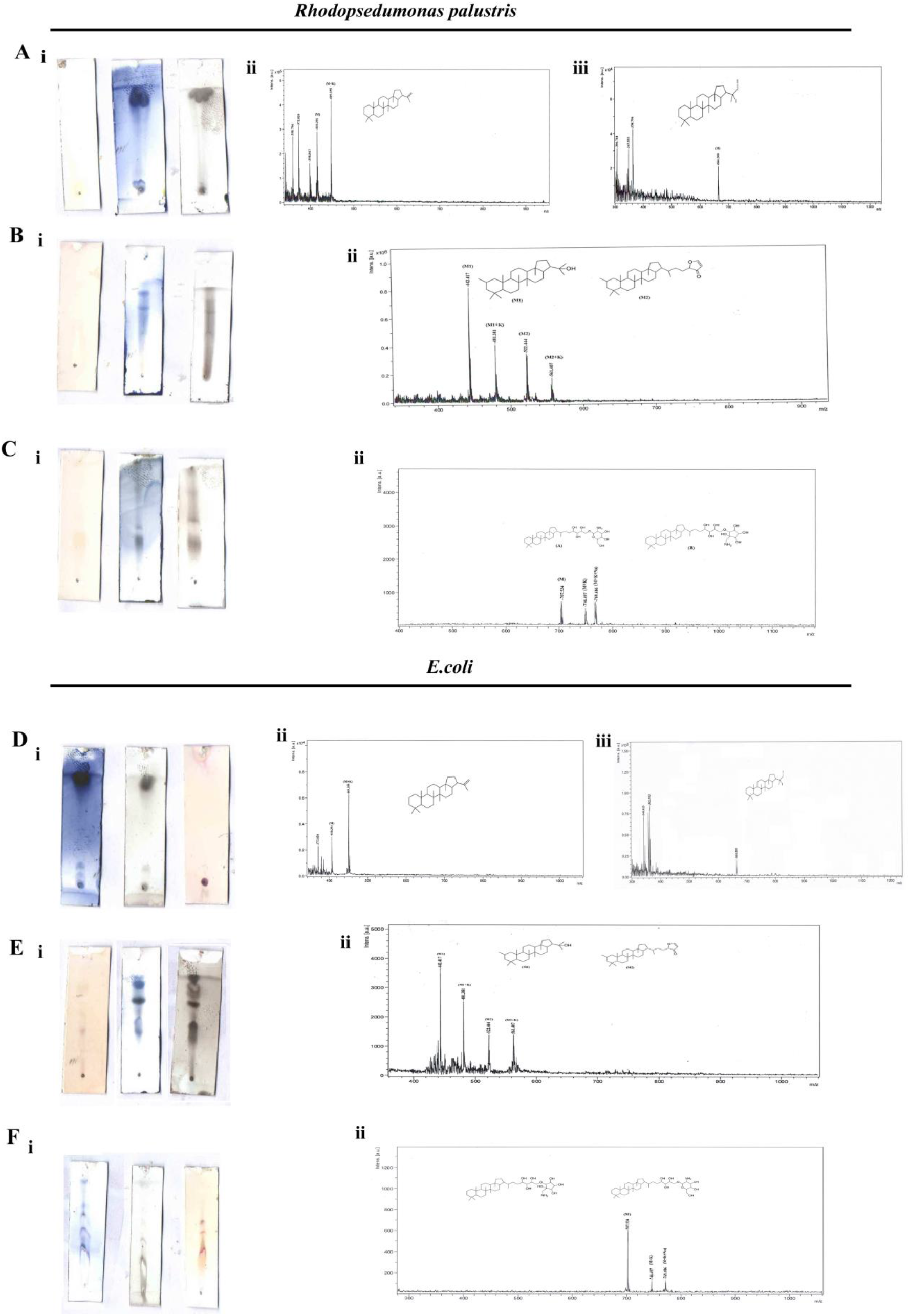
TLC and MALDI-MS of the fine purified probable Hopanoid (HOP) compounds. **(Ai, Bi, Ci)** TLC plates charred by three charring solutions likeNinhydrin, copper acetate, and Ammonium molybdate (from left to right) of different fractions of neutral lipid isolated from *Rhodopseudomnaspalustris* eluted in Hexane (1^st^ fraction) (A),30% Ethyl acetate-Hexane (2^nd^ fraction) (B), 5% Chloroform-Methanol (3^rd^ fraction) (C) following column chromatography. Amine group was detected by ninhydrin charring as indicated by Light brown color, nonpolar tail part was detected by copper acetate and Ammonium molybdate charring as indicated byblue and graycolor respectively. **(Aii, Bii, Cii)** MALDI-MS data profile of the fine purified compounds of different fractions of neutral lipids isolated from *Rhodopseudomnaspalustris* eluted in Hexane (A),30% Ethyl acetate-Hexane (B), 5% Chloroform-Methanol (C) following column chromatography was indicated by either molecular ion(M) peak or Na/K added molecular ion(M+Na/K) peak comparable with that of member of Hopanoid (HOP) family. **(Aiii)** MALDI-MS data profile of the fine purified 1^st^ fraction of neutral lipid isolated from *Rhodopseudomnaspalustris* eluted in Hexane following iodination to detect the degree of Unsaturation. **(Di, Ei, Fi)** TLC plates charred by three charring solutionslike Ninhydrin, copper acetate, and Ammonium molybdate of different fractions of neutral lipids isolated from *Escherichia coli* eluted in Hexane (1^st^ fraction) (D),30% Ethyl acetate-Hexane (2^nd^ fraction) (E), 5% Chloroform-Methanol (3^rd^ fraction) (F) following column chromatography. Amine group was detected by ninhydrin charring as indicated by Light brown color, nonpolar tail part was detected by copper acetate and Ammonium molybdate charring as indicated byblue and graycolor respectively. **(Dii, Eii, Fii)** MALDI-MS profile of the fine purified compounds of different fractions of neutral lipids isolated from *Escherichia coli* eluted in Hexane (A),30% Ethyl acetate-Hexane (B), 5% Chloroform-Methanol (C) following column chromatography was indicated by either molecular ion(M) peak or Na/K added molecular ion(M+Na/K) peak comparable with that of member ofHopanoid (HOP) family. **(Diii)** MALDI-MS data analysis of the fine purified 1^st^ fraction of neutral lipid isolated from *Escherichia coli* eluted in Hexane following iodination to detect the degree of Unsaturation.

**Figure 2.**
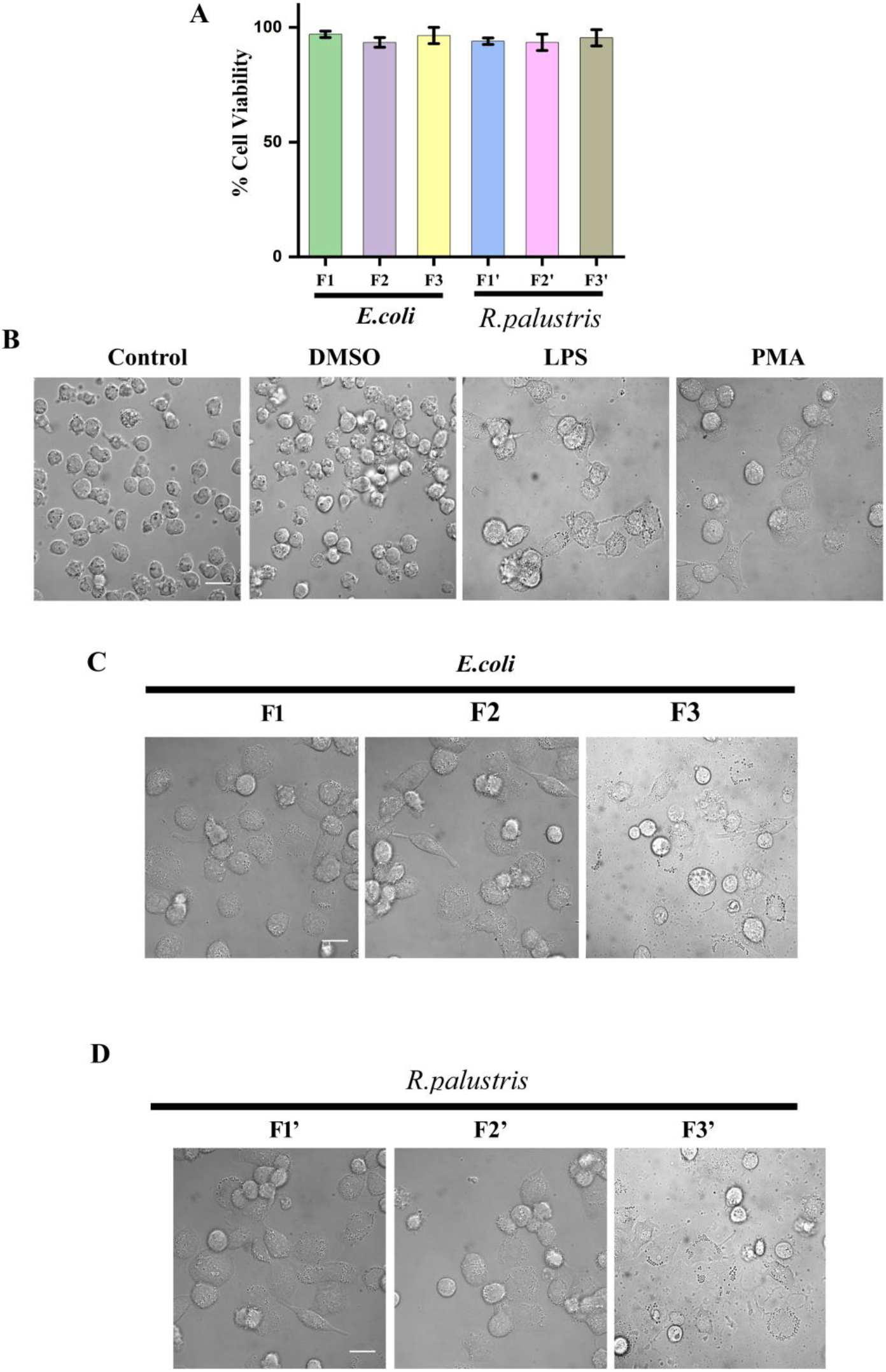
Cellular viability andmorphological analysis of monocytes post-treatment with probable Hopanoid (HOP) compounds. **(A)** Graphical quantification of cellular viability of monocytes (THP-1 cells) 48 hours post-treatment with the neutral lipid fractions isolated from *E*.*coli*(F1, F2, F3)and *Rhodopseudomonaspalustris* (F1’, F2’, F3’). **(B)** DIC images of control, DMSO(vehicle), LPS, and PMA stimulated THP-1 cells 4 days post-seeding. **(C-D)** DIC images of THP-1 cells stimulated with different neutral lipid fractions isolated from *E*.*coli*(C)and *Rhodopseudomonaspalustris* (D). Scale bar 15μm.Graphical analyses are performed with standard error bars after repeating the experiment at least thrice.

### Analysis of cellular viability and morphological features of monocytes post-treatment with probable Hopanoid compounds

The morphological difference is considered as a crucial parameter to distinguish between the monocytic and macrophagic status of the cell. To examine the immunomodulatory effects of the isolated neutral lipids (hopanoids),we have taken the well-established spherically shaped, suspension-cultured monocytic cell line THP-1 and HL-60, as our model system. Initially, we assayed the cellular viability of monocytes in response to bacterial hopanoids by MTT assay. No cytotoxic effects were observed even after 48 hours of treating the cells with the different fractions of neutral lipids compared to control (Fig. 2A). LPS and PMA are two widely used and well-established chemical stimuli, known to induce monocyte to macrophage differentiation. So in this study, LPS and PMA are used as positive controls to conclusively establish the cellular status of monocytes post-treatment with the probable hopanoid fractions. Fig.2C and D denote the cellular morphology of THP-1 cells post-treatment with probable hopanoids. In comparison with untreated control and DMSO (vehicle) treated sets (Fig.2B), the probable hopanoids treated cells adhered to the substratum and acquired heterogeneous morphology. Moreover, similar to LPS/PMA treatment hopanoids also resulted in the loss of circularity, an increase in the area, and the acquirement of elongated morphology in many cells (Fig.2B-D). To further verify our data, we did a similar experiment with the HL-60 cell line. Here, also we observed similar morphological alterations characteristic of differentiatedmacrophageinthe neutral lipid treated set (Fig.S3).

### Marker profiling of monocytes post-treatment with probable Hopanoid compounds

The differentiation of monocytes to macrophages is correlated with drastic changes in the expression of surface markers that are established signatures of these immune cells. Bone marrow-derived monocytes show elevated levels of CD35 and CD14 whereas fully functional tissue-resident macrophages display enhanced CD68 and CD64 on their cell surface(Fig.3Ai, Bi, C). We investigated the surface marker profile in THP-1 cells challenged with known immuno-stimulants LPS and PMA as well as with the probable hopanoid fractions isolated from *Escherichia coli* and *Rhodopseudomonaspalustris*. Semi-quantitative RT PCR and immunoblotting data reveal the distinct pattern of surface marker expression in the treated cells which show comparable similarity with those treated with LPS/PMA thereby confirming their ability to trigger differentiation(Fig.3A-B). Additional validation was done using immunostaining to confirm this pattern of marker alteration. The co-immunostained images depict the presence of CD35 (red fluorescence) in the undifferentiated cells and lack of CD68 expression. On the contrary, treated cells show a reverse pattern with maximal CD68 (green fluorescence) expression and complete absence of CD35 expression which indicates achievement of macrophage status of the cells(Fig.3D). Marker analysis in the HL-60 cell line has also validated our conclusion (Fig.S4).

**Figure 3.**
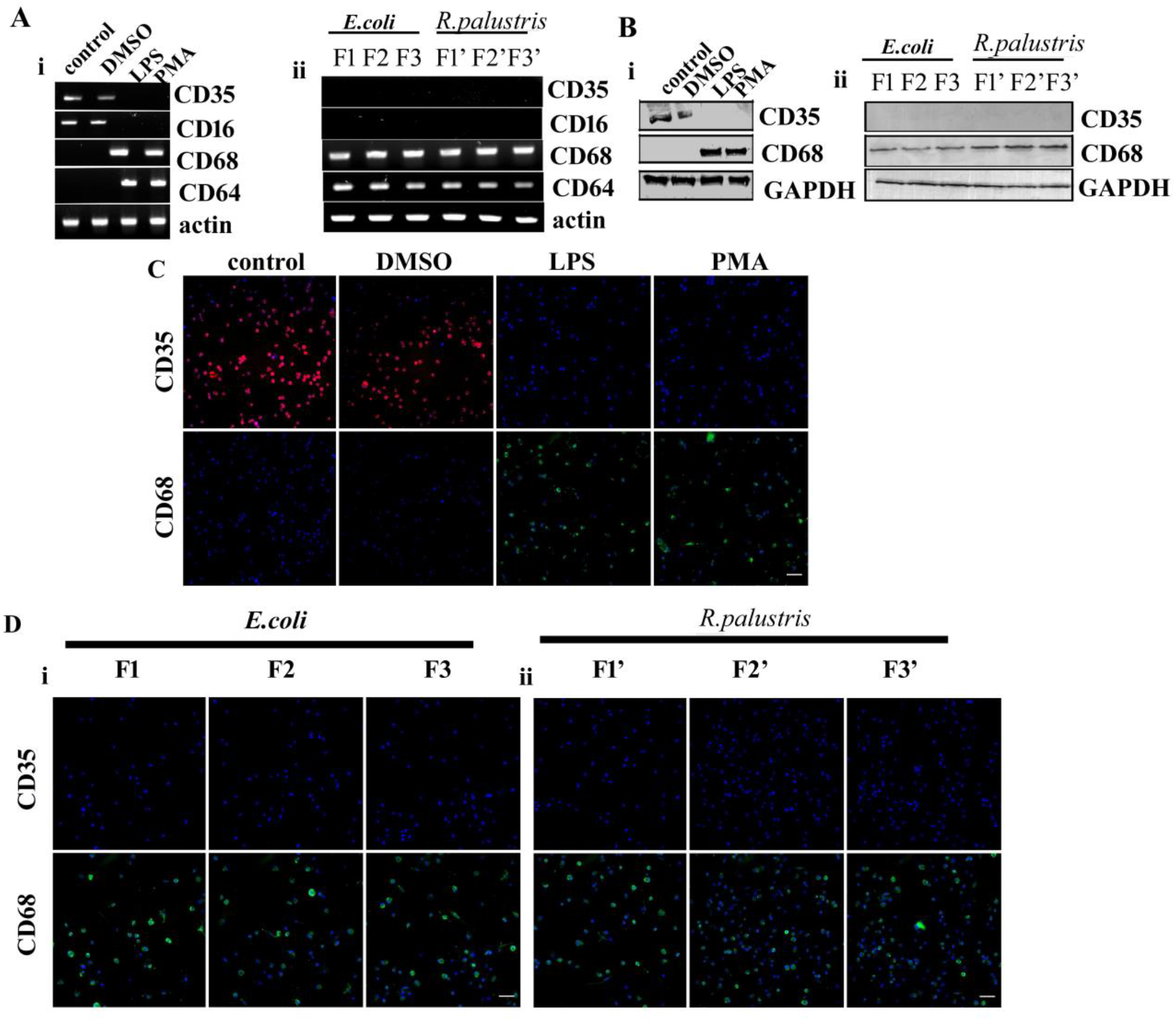
Marker profiling of monocytes post-treatment with probable Hopanoid (HOP) compounds. **(Ai)** Semi-quantitative RT PCR data of monocyte/macrophage markers viz. CD35, CD16, CD68, CD64 of THP-1 cells under untreated and treated conditions with DMSO, LPS or PMA. **(Aii)** Semi-quantitative RT PCR data of monocyte/macrophage markers of THP-1 cells stimulated with different neutral lipid fractions isolated from *E*.*coli*(first 3 lanes)and *Rhodopseudomonaspalustris*(4^th^-6^th^ lanes) 4 day post treatment. **(Bi)** Western Blot analysis of monocyte/macrophage markers viz CD35 and CD68 of THP-1 cells under untreated and treated conditions with DMSO, LPS or PMA. **(Bii)** Western Blot analysis of monocyte/macrophage markersof THP-1 cells stimulated with different neutral lipid fractions isolated from *E*.*coli*(first 3 lanes)and *Rhodopseudomonaspalustris*(4^th^-6^th^ lanes) 4 day post treatment. **(C)** Fluorescent microscopic images of CD35,CD68 immunolabelled and DAPI stained THP-1 cells under untreated and treated conditions with DMSO, LPS or PMA. **(D)** Fluorescent microscopic images of CD35,CD68 immunolabelled and DAPI stained THP-1 cellsstimulated with different neutral lipid fractions isolated from *E*.*coli*(first 3 panel)and *Rhodopseudomonas palustris* (last 3 panel). Scale bar 20μm.

### Qualitative estimation of intracellular organelles (lipid droplet, lysosome, and mitochondria) in monocytes post-stimulation with probable Hopanoid compound

Massive structural and biochemical modification occurs on-course of differentiation. Differentiation is an extremely energy-consuming process linked with significant changes in the mitochondrial and lipid droplet (LD) dynamics. The examination of the LD profile in the monocytes treated with the probable hopanoid fractions using BODIPY dye reveals the mass generation of lipid droplets compared to control (Fig.4A-B).The mitochondrial fission-fusion cycle and lysosomal proliferation were also monitored using MITOTRACKER and LYSOTRACKER dyes. Spectrofluorometric quantification of the stained cells showsa much higher intensity in the lipid-treated cells bearing resemblance to those cells challenged with LPS or PMA(Fig.4C-D).

**Figure 4.**
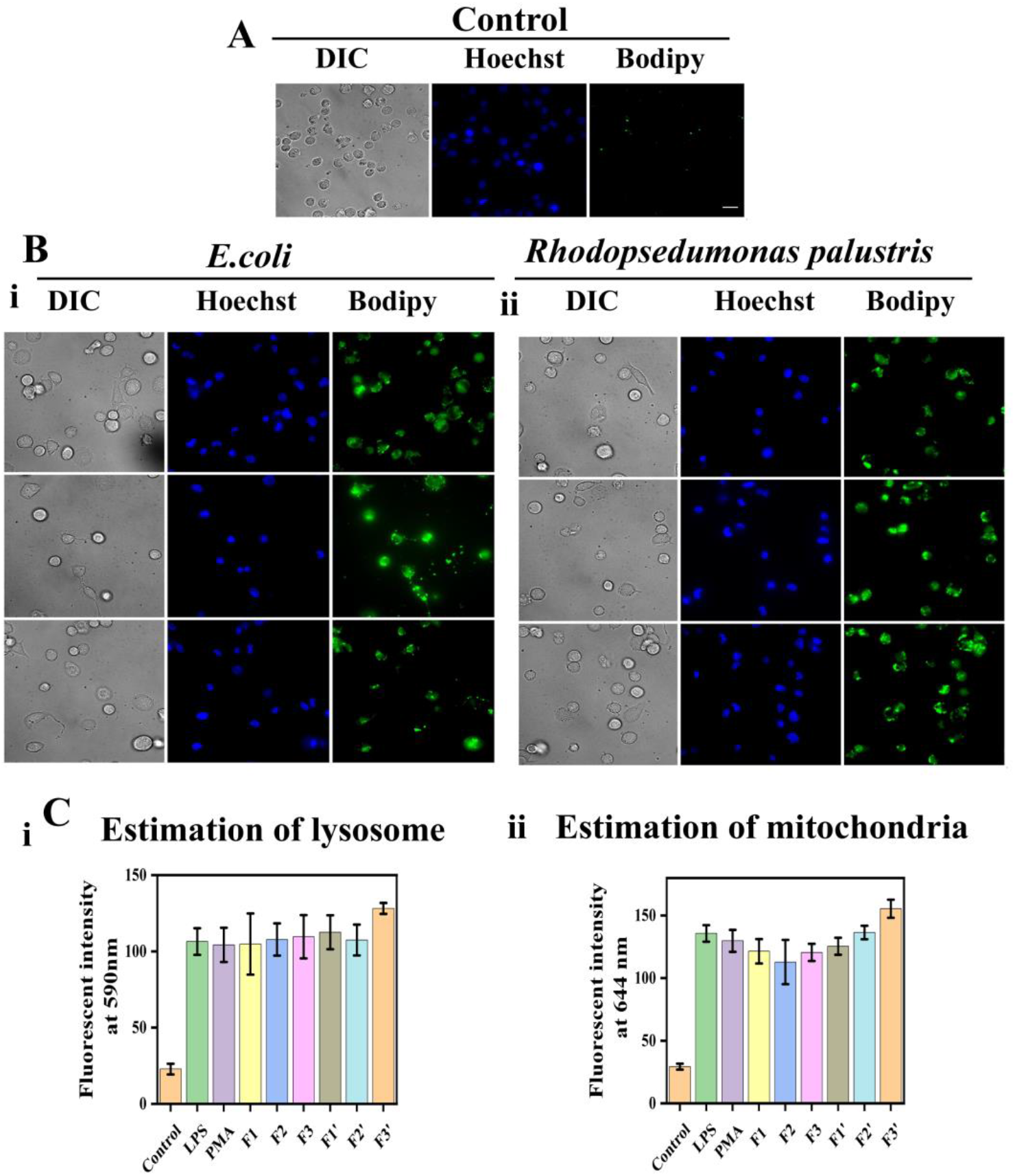
Qualitative estimation of intracellular organelles (lipid droplet, lysosome, and mitochondria) in monocytes post-stimulation withprobable Hopanoid (HOP) compounds. **(A-B)** DIC (left panel) and fluorescent images of Hoechst stained (middle panel) and Bodipy (right panel) stained THP-1cells unstimulated (A) and stimulated with different neutral lipid fractions isolated from *E*.*coli*(Bi)and *Rhodopseudomonaspalustris*(Bii). **(C)** Graphical analysis depicting fluorescent intensity of lysotracker (i) and mitotracker (ii) stained THP-1 cells stimulated with different neutral lipid fractions isolated from *E*.*coli*(F1, F2, F3) and *Rhodopseudomonaspalustris*(F1’, F2’, F3’). Scale Bar 10μm. Graphical analyses are performed with standard error bars after repeating the experiment at least thrice.

### Analysis of the functional attributes of monocytes post-stimulation with probable Hopanoid compounds

To further assess the effectiveness of the isolated neutral lipids (probable hopanoids) in inducing monocyte differentiation in-vitro, we resorted to analyze some of the established functional attributes of differentiated macrophages. Phagocytosis is one of the most designated properties of differentiated macrophages and we checked both the active and passive processes using RFP expressing *E*.*coli* and fluorescent polystyrene beads respectively. The percentages of cells undergoing phagocytosis treated with the three extracted lipid fractions are significant as compared to those treated with LPS or PMA(Fig.5A-B). Classically activated inflammatory macrophages secrete a portfolio of inflammatory cytokines which were evaluated by the semi-quantitative RT PCR method. The results reflect the transcriptional upregulation of inflammatory cytokines like IL-6, IL-8, and TNF-α in the treated groups with respect to untreated control and DMSO treated set(Fig.5C). Furthermore, macrophages are known to produce ROS and NO as a tool for killing engulfed pathogens. So we quantified the NO and ROS status among the different groups challenged with the isolated fractions and observations suggest that the hopanoids are at par with LPS/PMA to induce the generation of ROS and NO in the mononuclear cells(Fig.5D-E).

**Figure 5.**
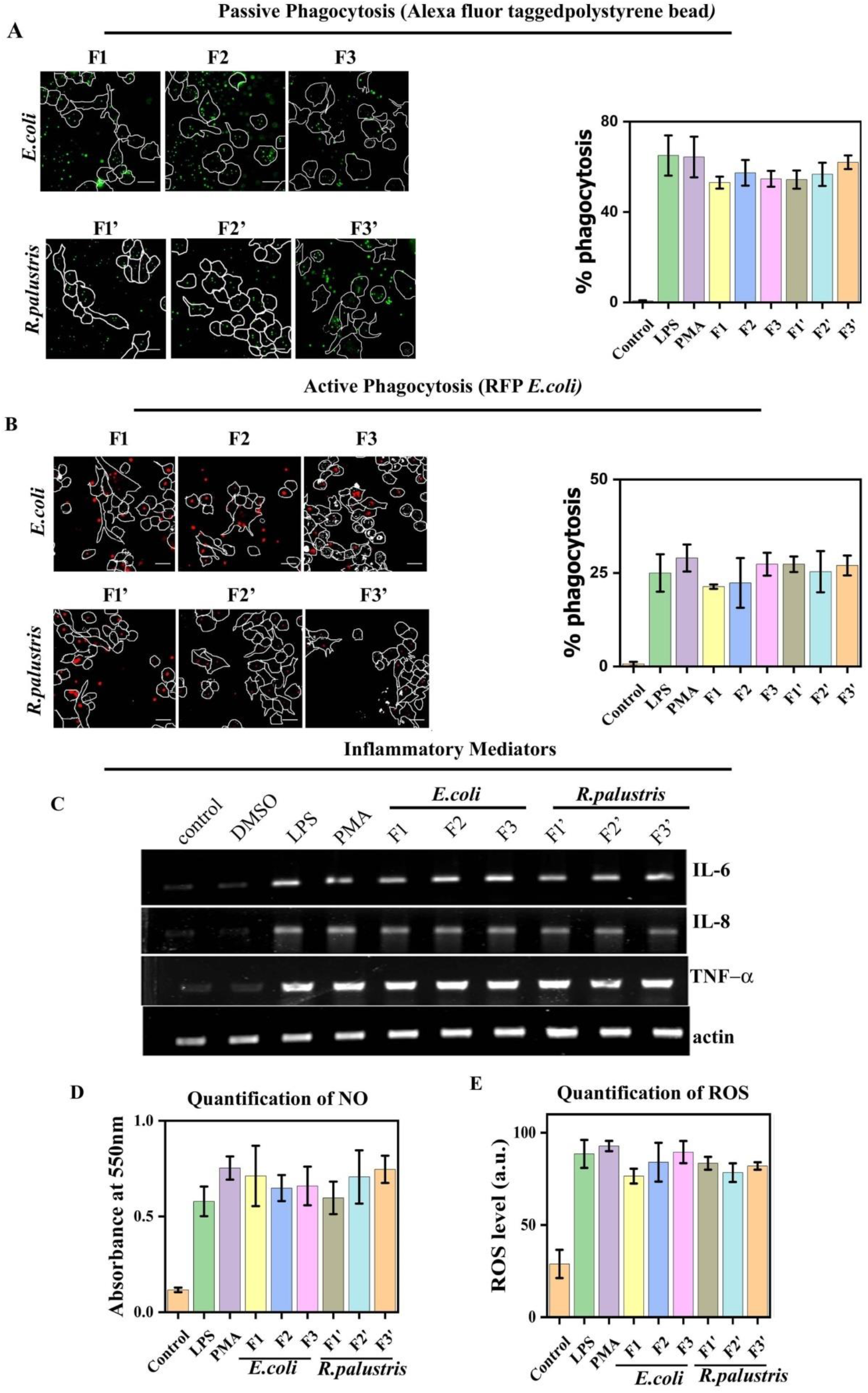
Analysis of the functional attributes of monocytes post-stimulation with probable Hopanoid (HOP) compounds. **(A)** Fluorescent images of passive phagocytosis (with fluorescent polystyrene bead)of THP-1cells stimulated with different neutral lipid fractions isolated from *E*.*coli* (F1, F2, F3) and *Rhodopseudomonas palustris* (F1’, F2’, F3’). Graphical representation of the percentage of passive phagocytosis is represented at the right. **(B)** Fluorescent images of active phagocytosis (with RFP expressing *E*.*coli*)of THP-1cells stimulated with different neutral lipid fractions isolated from *E*.*coli* (F1, F2, F3) and *Rhodopseudomonas palustris* (F1’, F2’, F3’).Graphical representation of the percentage of active phagocytosis is represented at the right. **(C)** Semi-quantitative RT PCR data of inflammatory cytokines viz. IL-6, IL-8, TNF-α of THP-1 cells stimulated with different neutral lipid fractions isolated from *E*.*coli*and *Rhodopseudomonas palustris*. Actin is used as control. **(D-E)** The quantitative comparison of NO (D) and ROS (E) production fromTHP-1 cells stimulated with different neutral lipid fractions isolated from *E*.*col i*and *Rhodopseudomonas palustris*. Scale Bar 10μm. Graphical analyses are performed with standard error bars after repeating the experiment at least thrice.

### Elucidation of signaling cascade involved in the monocyte to macrophage differentiation induced by the probable Hopanoid compounds

The differentiation process is ushered by the activation of a variety of signaling networks that command the various cellular changes. We along with other groups havealsoreported a plethora of signaling molecules contributing to the differentiation process notably MAP kinases and NF-κβ[45,46]. We investigated the involvement of these signaling molecules in the differentiation process induced by the probable hopanoids and our Western blot data suggests the occurrence of phosphorylation of P42/44, P38 MAP kinases as well as NFκβwhen treated with the probable hopanoidsvs the untreated control(Fig.6A). The quantification of these blots indicatesa manifold increase in the phosphorylation levels of these molecules resulting in the triggering of downstream signaling cascades(Fig.6A). To further reaffirm the involvement of these pathways, we used well-established inhibitors: U0126, SB203580 for MAP kinases, and NFκβI for NFκβ. The cells were pre-treated with the inhibitorsfollowed with stimulation with the isolated hopanoids fraction and subsequentlyco-immunostained with antibodies against CD35 and CD68 along with DAPI for nuclear staining. From the images in Fig.6B, we can visualize the absence of CD68 in the cells treated with the different fractions of probable hopanoidsisolated from the two Gram-negative bacterial strains highlighting the imperative role of these pathways in differentiation.Semi-quantitative RT PCR-basedtranscriptional profiling of the marker genes has also supported our postulation(Fig.6C).

**Figure 6.**
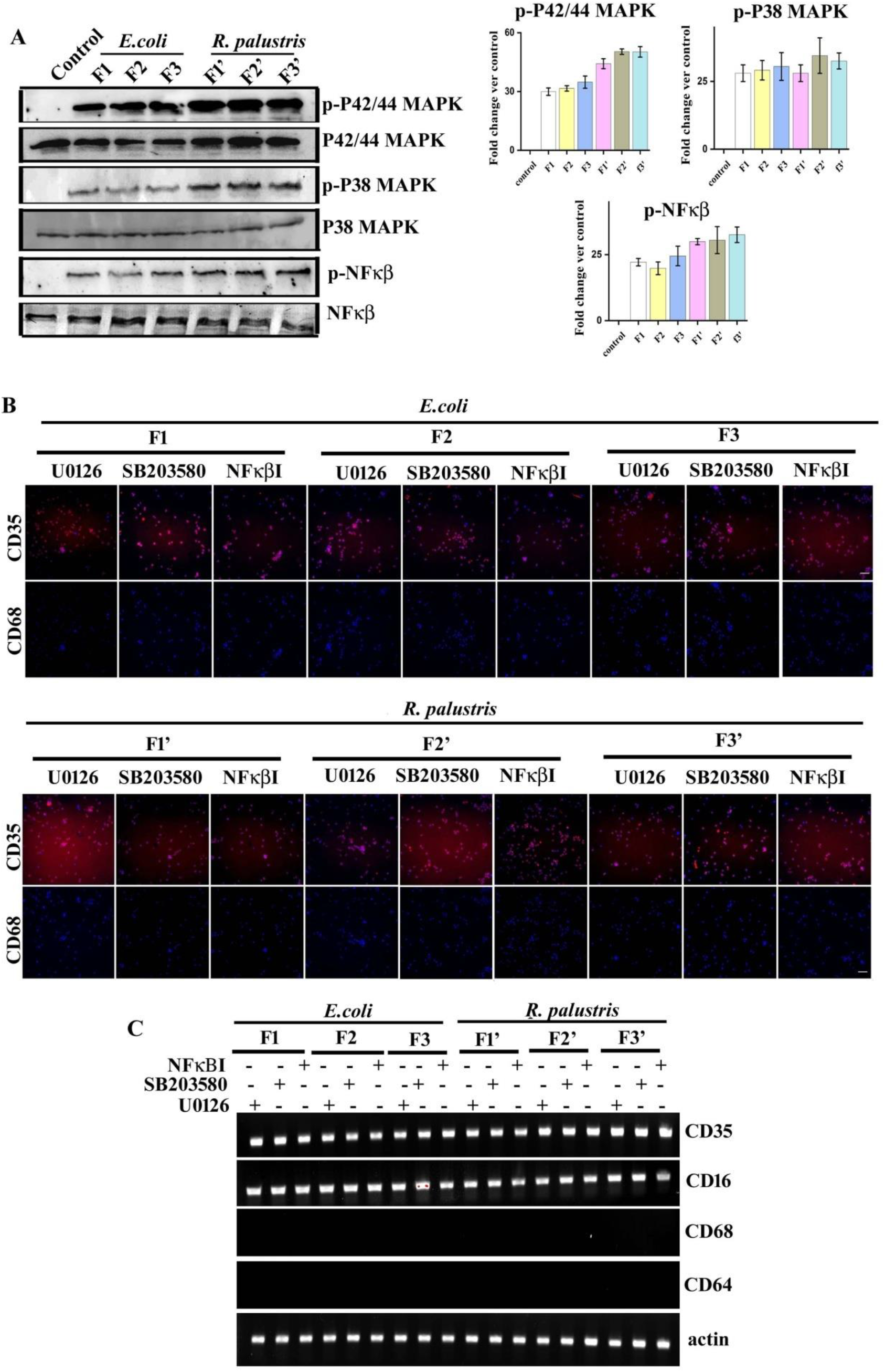
Elucidation of signalling cascade involved in the monocyte to macrophage differentiation induced by the probable Hopanoid (HOP) compounds. **(A)** Analysis of activation (phosphorylation) of P42/44MAPK, P38MAPK and NF-κβ of THP-1cells stimulated with different probablehopanoids fractions isolated from *E*.*coli*and *Rhodopseudomonaspalustris*via Western Blot. **(B)** CD 35 (upper panel) and CD 68 (lower panel) immunostained and DAPI stained images of U0126 (inhibitor of P42/44 MAPK), SB203580 (inhibitor of P38MAPK), NF-κβI (inhibitor of NF-κβ), treated THP-1 cells stimulated with 1^st^ fraction (left 3column), 2^nd^ fraction (middle 3column) and 3^rd^ fraction (Right 3 column) isolated from *E*.*coli* (Upper) and *Rhodopseudomonas palustris* (lower)4 days post seeding. **(C)** Semi quantitative RT PCR of CD 35, CD 16, CD 64 and CD 68 of U0126, SB203580,NF-κβI treated THP-1 cells 4 days post stimulation with different hopanoids fractions isolated from *E*.*coli*and *Rhodopseudomonaspalustris*.. Scale Bar 20μm.

### Autophagy is induced in monocytes in response to the treatment with probable Hopanoid compounds

Differentiation is a highly intricate process comprising of a nexus of cellular remodeling and organellar turnover. Autophagic recycling is a crucial mediator of differentiation and the advent of autophagy is often considered as an analytical marker of monocyte differentiation. We scanned for the presence of prominent autophagy markersmicrotubule-associated protein light chain 3 (LC3) to assay autophagy induction. Upon sensing the induction signal of autophagy, cytosolic diffused LC3 is processed to punctuate LC3 which characteristically represents the initiation of an autophagic response and confirms the subsequent formation of autophagosomes. During autophagy, cytosolic LC3-I is converted to its lipidated LC3-II form; and gets docked on newly forming autophagophores. In untreated monocytes, LC3 is present in a diffused pattern whereas, a significant increase in the LC3-IIB puncta was evident in stimulated cells (Fig.7A, S7A,C).Western blot data also affirmed the processing ofcellular LC3-IIB in monocytes in response to the probable bacterialhopanoids(Fig.7B, S7B). This finding was further corroborated by CYTO-ID staining(Fig.7C, S7E). CYTO-ID Autophagy Detection Kit labels theautophagic vacuoles and monitors autophagic flux in live cells using a novel dye that selectively labels accumulated autophagic vacuoles[47][48].Next, we evaluated the autophagy maturation in stimulatedmonocytes.Thefusion ofautophagosomes with lysosomes is termed autophagy maturation. So to assayautophagy maturation, we used DQ Red BSA, a fluorogenic substrate for acid-resistant proteases. Withinautophagolysosome, BODIPY tagged highly quenched form of DQ Red BSA is hydrolyzedleading to the formation of bright fluorescent products[49][50]. The fluorescence of DQ Red BSA strengthening our claim that bacterial hopanoid induces autophagy maturation with subsequent monocyte to macrophage differentiation(Fig.7D, S7G).

**Figure 7.**
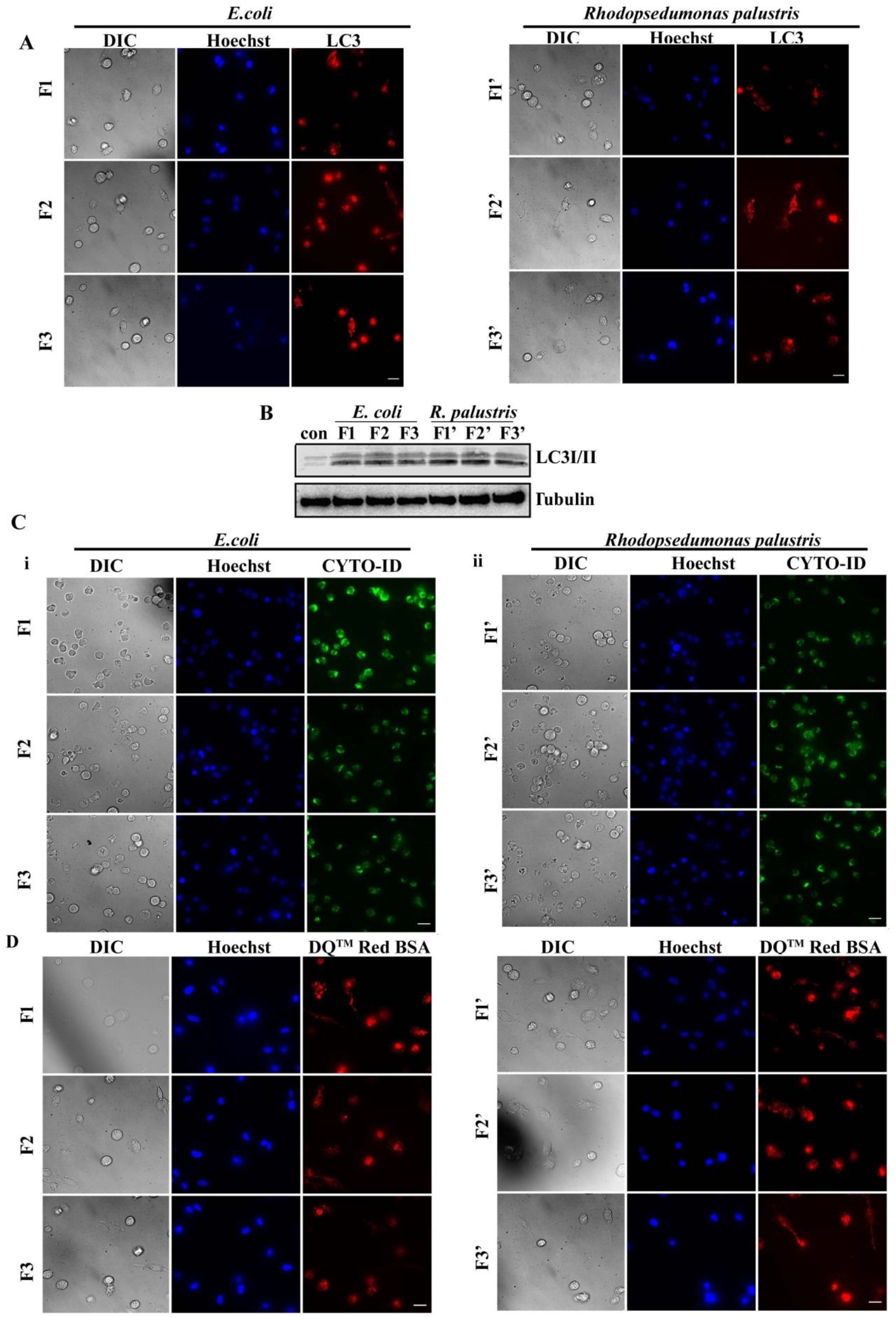
Autophagy is induced in monocytes in response to the treatment with probable Hopanoid (HOP) compounds. **(A)** DIC (left panel) and fluorescent images of Hoechst stained (middle panel) and LC3-IIB (right panel) immunostained THP-1cells stimulated with 1^st^fraction (upper), 2^nd^ fraction (middle) and 3^rd^ fraction (lower) of neutral lipid isolated from *E*.*coli*(left panel)and *Rhodopseudomonaspalustris* (right panel). **(B)** Western Blot of LC3 I/II in THP-1cells stimulated with 1^st^ fraction (upper), 2^nd^ fraction (middle) and 3^rd^ fraction (lower)of neutral lipid isolated from *E*.*coli*and *Rhodopseudomonaspalustris*. Tubulin is used as the loading control. **(C)** DIC (left panel) and fluorescent images of Hoechst (middle panel) and CYTO-ID (right panel) stained THP-1cells stimulated with 1^st^ fraction (upper), 2^nd^ fraction (middle) and 3^rd^ fraction (lower) of neutral lipid isolated from *E*.*coli*(i)and *Rhodopseudomonaspalustris*(ii). **(D)** DIC (left panel) and fluorescent images of Hoechst (middle panel) and DQ™ Red BSA (right panel) stained THP-1cells stimulated with 1^st^ fraction (upper), 2^nd^ fraction (middle) and 3^rd^ fraction (lower) of neutral lipid isolated from *E*.*coli*(i)and *Rhodopseudomonaspalustris*(ii).Scale Bar 20μm.

## Discussion

The mononuclear phagocytic system comprises blood circulating monocytes and tissue-resident macrophages[51]. Being a primary sentry of our defense system, monocytes and macrophages bridge innate immunity with adaptive immunity. Blood monocytes get recruited to the tissue and subsequently differentiate into mature macrophages[4]. A variety of stimuli is known to induce the monocyte to macrophage differentiation [52][53][54]. Different cytokines such as Macrophage-colony stimulating factor (M-CSF), Granulocyte monocyte-colony stimulating factor (GM-CSF), IL-1β, IFN-γ, etc are reported to persuade differentiation. Apart from cytokines, different hormones like growth hormone, leptin, insulin-like growth factor-1, steroid hormones, thyroid-stimulating hormone, prolactin, neurohypophyseal hormones, and T3-T4 hormone are also proclaimedto have an immune modulatory effect[55][56][57][58]. On the other hand, steroid-based Vitamin D_3_ {1,2 5-dihydroxy vitamin D_3_} is a potent inducer of monocyte to macrophage differentiation [59]. Apart from these, different pathogens like bacteria, viruses,etc [60][61]as a whole or some of their cellular remnants are also considered as activatorsof monocytes. For example, LPS (lipopolysaccharide) present in the outer membrane of Gram-negative bacteria [52],lipoteichoic acid, muramyl dipeptide, formylmethionine-leucine-phenylalanine, protein A, D-mannose are described to take part in the immune modulation of the host.

Although several discrete studies have been performed to isolate, identify, and characterize biochemical properties of hopanoids,the effect of bacterial hopanoids on the host immune system is not well reported. Interestingly, the total non-polar lipid of Gram-negative bacteria was able to induce monocyte to macrophage differentiation. This further affirms our hypothesis that apart from LPS, some other membrane components of Gram-negative bacteriacan exhibit the immune-modulatory effect. Thus, in the present manuscript,weexplored whether bacterial hopanoidsinfluence human monocytes by analyzing and comparing their characteristicmorphological parameters. Moreover, we have considered the gain of other structural as well as functional attributes associated with thedifferentiation process to analyze the cellular state of these stimulated cellsand compared them with those of macrophages stimulated by LPS and PMA.

In the recent past, several groups have standardized the basic protocol for hopanoids isolation from bacteria like *Rhodopseudomanospalustris*. Here, we were also able to successfully isolate severalactive compoundsin the neutral lipid fractions of*Rhodopseudomonaspalustris* and we speculated them as probable hopanoid. Moreover, the hopanoid isolation protocol from other Gram-negative bacterialike *Escherichia coli*is not reported. In this work, we have applied the same isolation protocol reported for *Rhodopseudomonaspalustris*for extraction of different neutral lipid fractions from *Escherichia coli*, and fortunately, we could detect the presence of a similar compound having a comparable molecular weight in the neutral lipid fractions of *Escherichia coli*. Following isolation, we have structurally and biochemically characterized those active compounds and due to the similarities with the reported hopanoids, we have considered that compound 1 (Diploptene), compound 2 and compound 3 (2 methyl diploterol) may be present in the isolated neutral lipid fractions. Incidentally, hopanoidswerenot found in the neutral lipid fractions isolated from the two typical Gram-positive organisms like *Staphylococcus aureus*and *Bacillus subtilis*.

In this manuscript, we have shown that like LPS/PMA, the probable bacterial hopanoids arealso capable of inducing similar morphologicalalterations in human monocytes ubiquitously. Although the probable hopanoids treatment resulted in a typical change of cellular morphology, mere morphological differences should not be considered as an ultimate confirmation of any altered cellular state. So the use ofstage-specific protein markers has been introduced to identify the cellular status of monocyte or macrophage. From the literature survey,we have chosen CD 35 and CD 16 as monocytic, CD 64, and CD 68 asmacrophagic markers [62][63][64][65][64][66]. In this manuscript, we have demonstrated that similar to LPS/PMA, the treatment of speculated hopanoids in monocytes leads to a significant increase in expression of macrophage-specific marker along with simultaneous loss of monocyte specific marker;suggesting an immune-modulatory property of bacterial hopanoids in the context of monocyte-macrophage differentiation.

The digestive function of macrophage is powered by lysosomal hydrolases for the degradation of ingested particles[67]. Consistent with the heightened degradative capacity by the virtue of greater numbers oflysosomes, simultaneous higher energy production is also requiredby an increased number of mitochondria.Several reports suggest the accretion inthe number of lysosomes and mitochondria is associated with monocyte to macrophage differentiation [68][69]. Here also, we identified an escalation in the numbers of mitochondria and lysosomes in monocytes treated with the probable hopanoids in comparison with untreated control. In the recent past, Ruiz et al and Hartigh et al have shown differential expression of lipid metabolism associated genes in monocytes and differentiated macrophages[70][71]. Thus lipid droplet accumulation is considered to be associated with monocyte to macrophage differentiation.In this manuscript, we have also found lipid droplet accumulation in hopanoid treated cells which can be considered as a hallmark of monocyte differentiation.

Apart from visual differences, stage-specific functional property like augmented phagocytosis enhanced inflammatory cytokine production, and elevated ROS and NO productionare also considered as characteristic properties of macrophages that distinguishes them from monocytes. As the name suggests, “Macrophages” are professional phagocytes and efficientlyremove dead cells and cellular debris.They are principally involved in the clearance of cellular debris during physiological homeostasis as well as in the removal of exogenous particles, including micro-organisms[4].ÉlieMetchnikoff has described the phagocytotic potential of macrophages as the key to immunity[72]. Here, we demonstrate thatin*in-vitro* culture as well,the probable bacterial hopanoids have elicited similar phagocytotic property in comparison to untreated monocytes similar to that of LPS/PMA.

Duque et al have reported that through the production of an array of cytokines,macrophagesplay the central role as sentinels of the innate immune system and also mediate the transition from innate to adaptive immunity[16].Chomarat et al have shown that MCSF induced monocyte to macrophage differentiation is accompanied by the production of enhanced inflammatory cytokines[73]. Many other independent reports described that other inducers of monocyte differentiation also led to a similar increase in inflammatory cytokines production[74].Through this study, we have shown that the probable bacterial hopanoidscan promotesimilar enhancement of inflammatory cytokine production in the context of monocyte to macrophage differentiation. During inflammation, macrophages secrete several soluble and diffusible inflammatory mediators which are active both locally (at the site of tissue damage and infection) and at more distant sites. In the recent past,McNeill et al have reported that i-NOS gene expression is found to enhanceseveral folds in treated THP-1 cells compared to control[14]. As a result of enhanced accumulation of i-NOS gene products in macrophages, these cells also secrete nitric oxide in the supernatant. Sekhar et al have also shown that chemical stimulation has led to a significant increase of NO secretion by the monocytes [17]. In agreement with these reports, in this study, we also have shown an enhanced production of nitric oxide in response to probable hopanoids treatment.

Reactive oxygen species (ROS) are key signaling molecules that play an important role in the progression of inflammation. Kuwabara et alhave shown an enhanced ROS generation by macrophages at the site of tissue damage mediate the inflammatory response and aid in the ailment of damaged tissues[75]. So aggravation of cellular ROS accumulation in macrophages compared to monocytes can also be accounted as a parameter of differentiation.Here, we demonstrate a similar escalation of ROS production by the cells treated with the probable hopanoids.

Our previous publications depict that the MAPK-NFκβ signaling pathway is a crucial regulator of monocyte to macrophage differentiation[45,46].Commensuratewith other reports, we have also shown that inhibition of the activation of this signaling cascade abates differentiation.In this manuscript, our results suggest that hopanoids treatment also leads to the activation of MAPK-NFκβ signaling pathway for the induction of monocyte to macrophage differentiation.

Another imperativefacet of monocyte to macrophage differentiation is autophagy. Zhang et al and Jacquelet al have shown autophagy to be intrinsically associated with the monocyte differentiation process[12,13][76].Many independent reports described that the chemical inducer like GMCSF, MCSF, PMA, Vitamin D3 [77][78]induces autophagy to persuade monocyte to macrophage differentiation process. Here, we demonstrate the active participation of autophagy in the probable hopanoids-induced monocyte differentiation process.

Macrophages are a major regulator of our immune system. As a part of innate immunity, they primarily engulf the intruding foreign pathogen, digest them, secrete cytokines and other soluble biochemicals to eliminate the dying damaged cells and also aid in the process of tissue repair[79][4].Macrophages also act as antigen-presenting cells (APC) and remain involved in the elicitation of adaptive immunity. Thus, it is crucialtosearch for new immuno-modulators to facilitate monocyte to macrophage differentiation.In this manuscript, we have convincingly established that along with LPS, Gram-negative bacteria also modulate the host immune system through hopanoids. In a nutshell, this study showed that bacterial hopanoids can induce cellularbiochemical changes in vitro associated with modulationof monocyte phenotype to promote their differentiation to macrophages.These studies will be useful to design better therapeutics against bacterial infection with minimal side effects.

## Declaration of the competing interest

All authors declare no conflict of interest.

## Funding

This work is supported by Department of Science and Technology, India [grant no. SR/SO/BB-0125/2012]

## Acknowledgements

We acknowledge CSIR fellowship to AB (SPM), SM (SPM) and AG. We acknowledge Dr. Deepak Sinha for the microscope facility.

## Notes

### Competing Interest Statement

The authors have declared no competing interest.

